# Nanoscopic Stoichiometry and Single-Molecule Counting

**DOI:** 10.1101/524801

**Authors:** Daniel Nino, Daniel Djayakarsana, Joshua N. Milstein

## Abstract

Single-molecule localization microscopy (SMLM) has the potential to revolutionize proteomic and genomic analyses by providing information on the number and stoichiometry of proteins or nucleic acids aggregating at spatial scales below the diffraction limit of light. Here we present a method for molecular counting with SMLM built upon the exponentially distributed blinking statistics of photoswitchable fluorophores, with a focus on organic dyes. We provide a practical guide to molecular counting, highlighting many of the challenges and pitfalls, by benchmarking the method on fluorescently labeled, surface mounted DNA origami grids. The accuracy of the results illustrates SMLM’s utility for optical ‘-omics’ analysis.

## 1. Introduction

Recent advances in optical imaging have been so dramatic that light microscopy can now be used to directly quantify protein or nucleic acid numbers, modifications and interactions, replacing less sensitive biochemical techniques like enzyme-linked immunosorbent assays (ELISA) or colorimetric assays. This has given rise to the new fields of both ‘optical proteomics’ and ‘optical genomics’^[1]^. Already, these optical ‘-omics’ techniques have been employed to quantify amino acid content in short peptides and proteins^[2;3]^, perform system-wide analyses of protein and mRNA copy numbers in *E. coli* to better understand stochastic gene expression^[4]^, and probe complex protein networks involved in cancer formation^[5]^. With the advent of single-molecule localization microscopy (SMLM), which provides sub-diffraction limited images extending light microscopy from the microscopic to the nanoscopic^[6;7]^, there is the potential to significantly accelerate the optical ‘-omics’ fields, leading to the development of new platforms for the high-throughput characterization and identification of proteins and nucleic acids at ultra-low concentrations *in vitro* and in single-cells^[8]^.

The increased information content of single-molecule surements has significant advantages compared to bulk measurements. Additional parameters become accessible, like receptor occupancies, ligand density, protein subunit or complex stoichiometry, and so on. Moreover, when combined with the enhanced imaging resolution of SMLM (~ 10 nm lateral resolution, with slightly poorer depth resolution), spatial details such as local protein concentrations or ‘spatial transcriptomics’^[9]^ become available.

SMLM capitalizes upon the fluorescence intermittency (or ‘blinking’) that certain fluorophores may be induced to exhibit^[10;11]^. The strategy is to locate the central coordinates of individual fluorophores that are made to blink such that the odds of overlap between the fluorophores at any time is low. In this way, the fluorophores are effectively separated in time and a table of coordinates or ‘localizations’ is assembled from large stacks of sparsely imaged, single molecules. Note, a single blink may give rise to multiple localizations dependent upon the duration of the blink (or on-time) and the camera frame rate. Regardless, this localization table is essentially a register of every molecule imaged, implying that SMLM may alternatively be used as a molecular counting method, providing quantitative information on protein or nucleic acid abundance and stoichiometry. And unlike standard fluorescence counting techniques, such as extracting molecule number from a calibrated total intensity or from photobleaching steps^[12;13]^, counts may be obtained for clusters or aggregates of biomolecules separated by distances well below the diffraction limit.

Unfortunately, in practice there isn’t a simple one-to-one relationship between the number of localizations and the number of molecules. Most dyes and fluorescent proteins used in SMLM are known to cycle between photoswitchable states with the same fluorophore blinking several times before photobleaching, often resulting in an over-estimation of the number of molecules present^[14;15;16]^. A number of groups have devised approaches to molecular counting with SMLM that attempt to mitigate the error from repeated blinking. These methods often require detailed knowledge of the photophysical kinetics or utilize complex statistical techniques, such as hidden^[17]^ or aggregated Markov models^[18]^, to infer transition rates. For fluorophores with multiple dark states, multiple parameters need to be measured and/or inferred. A simpler, alternative approach is to measure the distribution of the number of blinks or localizations emitted by an ensemble of clusters. By relating the observed distribution to that obtained from a single fluorophore, one may quantify the number and stoichiometry of the clusters^[19;20]^.

Our molecular counting method, developed in Nino *et al.* ^[21]^, falls into this latter category and is built upon the common, empirical observation of an exponential number of blinks emitted from a single fluorophore ^[22]^. Here, we reformulate the theory developed in Nino *et al.* ^[21]^ so it’s relevant to molecular counting experiments with organic fluorophores imaged via the SMLM technique direct stochastic optical reconstruction microscopy (dSTORM)^[23]^. We then provide a practical guide to applying this theory to a molecular counting experiment, to quantify nanoscopic aggregates or clusters, and carefully benchmark the method in a controlled system of surface mounted DNA origami grids.

## 2 Molecular Counting

### 2.1 Estimating Molecule Number and Stoichiometry

We previously reported a closed-form, maximum-likelihood (ML) estimate for the number of molecules inferred from counting blinks in an SMLM measurement^[21]^. We also provided an analytic expression for the associated error on this estimate. For the case of dSTORM considered here, we must modify those relations. We assume that every dye blinks at least once since all the dyes are initially in a fluorescent state, and the vast majority of the dyes will cycle through a radical dark-state at least once before photobleaching. single-dye, *N* = 1, the probability distribution function for observing *B* ≥ 1 blinks takes the following form:

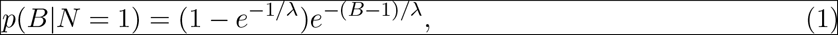

where λ is the characteristic number of blinks parametrizing the distribution. Note, Equation (1) differs from the form of the geometric distribution we utilized previously^[21]^ where we assumed that the distribution spanned the range *B* ≥ 0. For the case of *N* ≥ 1 dyes, the blinking distribution may be extended as follows:

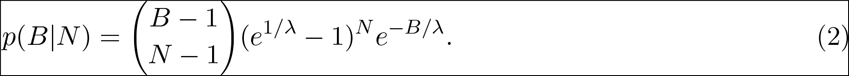

Following the previously detailed procedure^[21]^, with a binomial distribution modeling the labeling statistics of the form 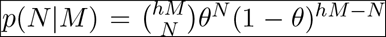 (*h* is the maximum number of labeling sites, *M* is the number of molecules, and *θ* is the fractional occupancy ranging from 0 to 1), the adapted formula for the ML estimate of the number of molecules given an observed number of blinks *B* is

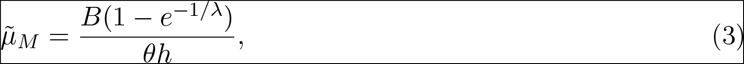

with an associated variance in the estimate of:

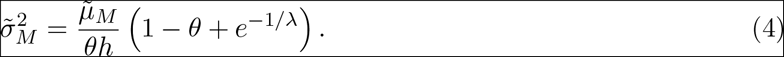

A similar set of analytical relations may be found assuming Poisson distributed labeling (see *Supporting Information*). Alternatively, if we have *a priori* knowledge of the number of molecules, we can make a maximum-likelihood prediction of the fractional occupancy *θ* inferred from Equation (3) as follows:

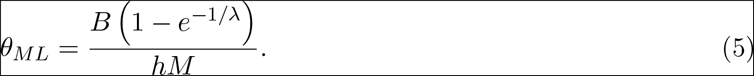

### 2.2 ML Calibration of Photophysics

Our method relies on quantifying certain photophysical properties of the photoswitchable dye employed, namely, the characteristic number of blinks λ. The brute force approach would consist of building up a histogram of the number of blinks per single fluorophore and then extracting λ by a non-linear least squares fit. This, however, requires larger sample sizes than extracting λ from a fit of the cumulative probability distribution. Another alternative is to simply count the number of blinks emitted by a known number of single-fluorophores, and then perform an ML estimate from the relation:

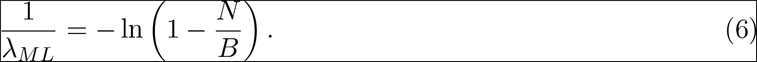

To calibrate λ one could sample sparsely labeled dyes such that each dye is individually resolved. This calibration could be an *in vitro* sample of organic dyes functionalized to a glass coverslip. Or *in vitro*, for example, it may be a protein containing only a single fluorescent label, known not to aggregate, and expressed at low enough levels that individual proteins may be resolved. What is essential is that the calibration reflects the same photophysics as when the fluorophores are bound to the actual target of interest. Admittedly, one must be careful when performing this calibration as the dye photophysics is often rather sensitive to its local environment^[24]^, as we will see.

### 2.3 The Thresholding Problem

A prerequisite to utilizing the approach presented here is that one can accurately tabulate the number of blinks in an SMLM measurement. In practice, this information is extracted from a localization table, which is a list of coordinates for each dye centre, often with an associated intensity of the dye’s point-spread function (PSF). The intensity is included to screen erroneous localizations detected by a localization algorithm arising from background noise, so a threshold is often imposed as a signal filter. Setting the threshold too high will discard many actual localization events, while setting too low a threshold will accept many spurious localizations.

In practice, the appropriate choice for this threshold is somewhat arbitrary and almost never reported in the literature. While a reasonable range of thresholds may result in an acceptable image reconstruction, for quantitative molecular counting applications, the choice of threshold can have a significant effect on the estimated count. Previously, we showed that the total number of blinks can be estimated in a threshold independent manner by extrapolating to zero threshold^[21]^. Here we clarify why such an approach is possible, which is mainly due to the temporal binning by the detection camera.

Figure 1a displays a time trace for the intensities of a blinking dye in the ideal case of constant fluorescence emission intensity. For sufficiently long lived dark-states, the long off-times mean that two independent blinks are unlikely to overlap within a frame. And for an average on-time of a few frames, which is common, a dye will rarely turn on and off within a single frame. With these two points in mind, we categorize each frame during a blink of a dye as either a ‘middle’ frame (if the dye is on in the neighbouring frame both before and after the considered frame) or as an ‘edge’ frame (if the dye either turns on or off within the frame).

**Figure 1.**
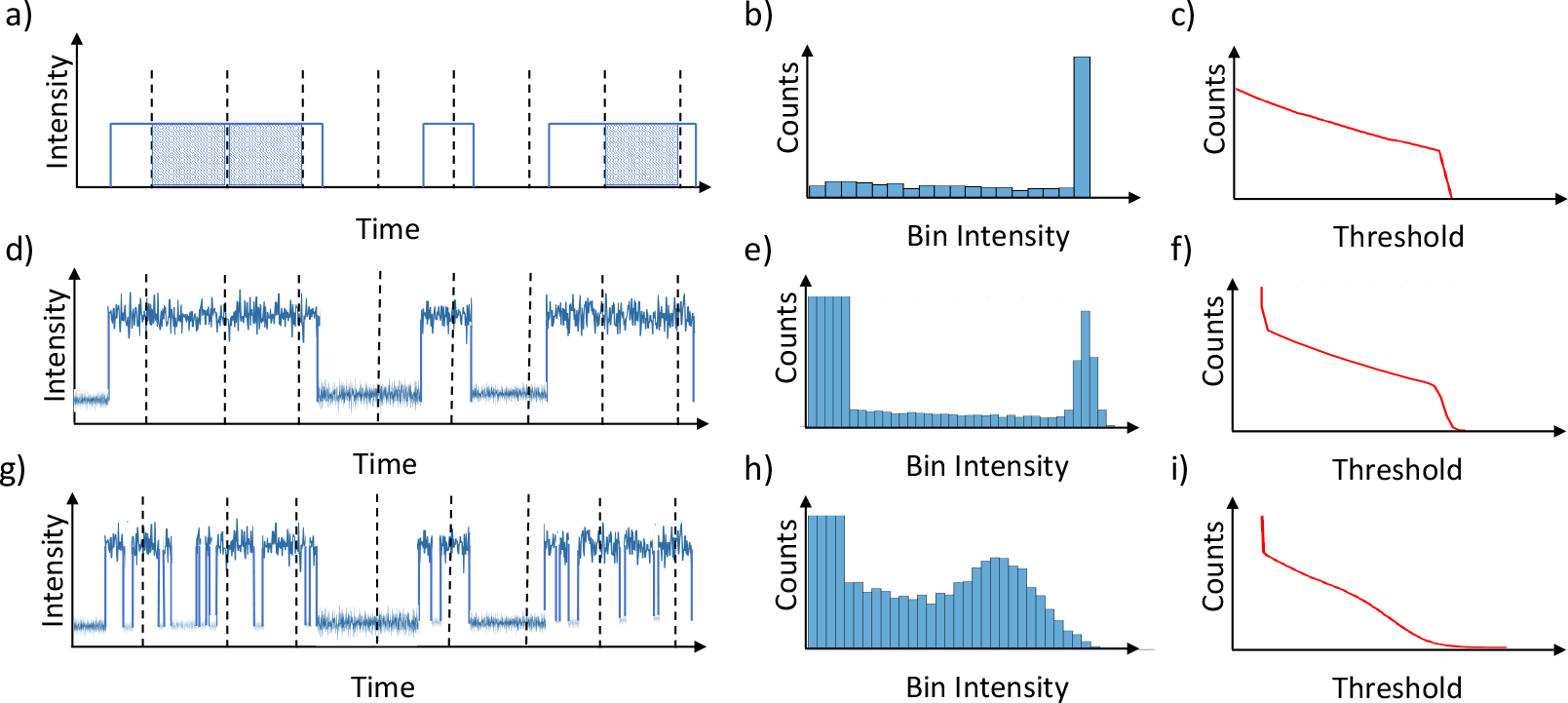
a) Ideal time trace of fluorescence intensity for a blinking dye where the on-state emits a constant number of photons per unit time. Dashed lines indicate discrete, camera frames, with ‘middle’ frames shaded, bordered by ‘edge’ frames. b) Localization counts vs. binned intensity. c) Localization counts vs. intensity threshold. d-f) Same as a-c, but for fluorescence displaying shot, detector, and background noise. g-h) Same as d-f, with the addition of triplet state blinking and/or excited singlet state quenching.

In an ideal SMLM measurement, for each frame that the dye is on, one would obtain a single localization. Figure 1b represents a histogram of measured intensity for each localization assuming the simplified emission dynamics of Figure 1a. The middle frames contribute the large peak at the intensity maximum, corresponding to dyes that are on for a full frame, while the edge frames produce a wide distribution of lower intensities. This broad distribution is essentially flat since since the moment at which a dye turns on or off within the frame is completely random, and the resulting intensities will range from zero to the full frame intensity with roughly constant probabilities. Now if we were to integrate this histogram starting at a given intensity threshold, we would generate a plot of the number of localizations vs. threshold that would display a broad, approximately linear region. As we have previously shown^[21]^, this linear region can be extrapolated to zero threshold yielding an estimate for the total number of localizations, even those hidden within the noise.

In reality, the fluorescence intensity is never constant, but fluctuates due to photon shot-noise, background and detector noise, etc. (Figure 1d). This noise has two effects. First, at low intensities, it gives rise to many false localizations, justifying the use of a threshold. Second, it broadens the peak at the high edge of the intensity distribution (Figure 1e). For sufficient signal-to-noise ratios, a relatively linear trend will still persist in a plot of the detected number of localizations vs. threshold (Figures 1f). However, if significant triplet state blinking and/or excited singlet state quenching is observed, for instance, the middle distribution can both broaden and shift toward lower intensities, and the extent of the linear region will be reduced (Figures 1g-i), or may disappear altogether.

## 3. Results and Discussion

To benchmark the SMLM counting technique presented here, we analyzed a system of surface mounted DNA origami grids. We first employ these nanostructures to calibrate for both the characteristic number of blinks λ emitted by the dyes and for the fractional occupancy of labeling *θ*. We then test the accuracy of our approach by directly imaging individual nanostructures and comparing to the ML estimate from the observed number of blinks.

### 3.1 Calibration of λ

As an illustration of the challenges to accurately obtaining the characteristic number of blinks λ, we first perform this calibration by SMLM imaging a system of spatially separated, DNA origami grids. The grids were each labeled with a single Alexa-647 dye centered on the grid. Results for this calibration, acquired at a low threshold (~ 650 photons, *N* = 1059 dyes), are presented in Figure 2. The insert to Figure 2a shows the characteristic, exponential dependency of the number of blinks obtained from a single dye. An exponential fit to this distribution, *f* (*B*)= (1 − *e*^−1/λ^)*e*^−(*B*−1)/λ^, yields λ = 3.6 ± 0.2 (*R*^2^ = 0.999). However, a preferred way to extract λ from the data, requiring less sampling, is to instead fit the cumulative distribution function *D*(*B*) = 1 − e^−B/λ^. From the same data, we find a slightly smaller and better constrained value of λ = 3.45 ± 0.05 (*R*^2^ = 0.995).

**Figure 2.**
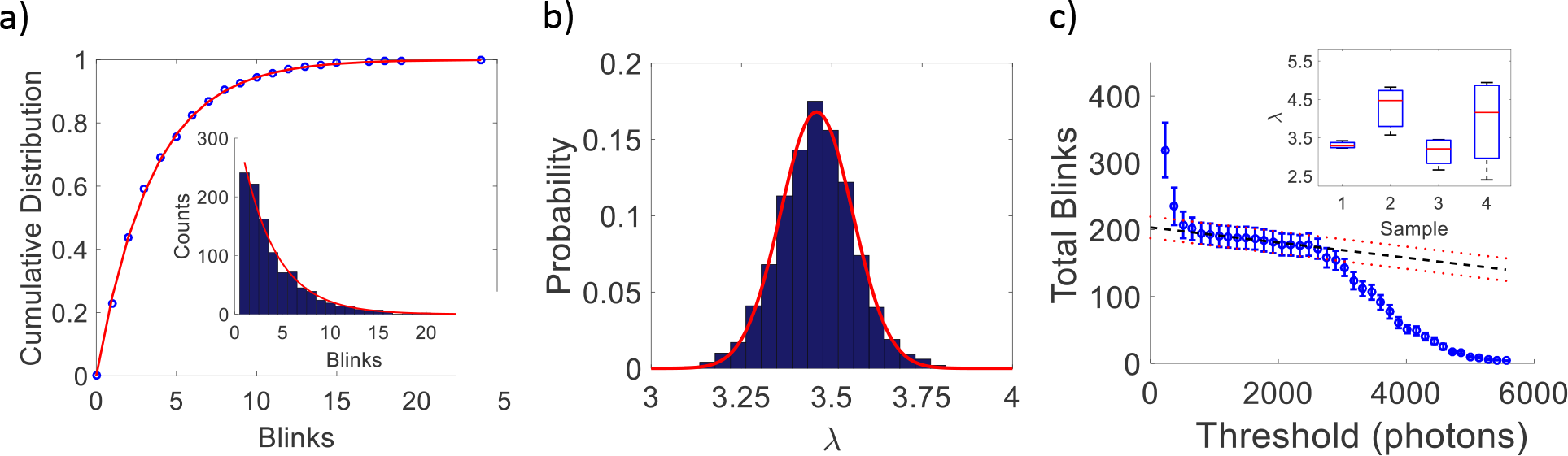
Estimating λ at low threshold (~ 650 photons, *N* = 1059 dyes) from: a) cumulative distribution function (λ = 3.45 ± 0.05, *R*^2^ = 0.995), inset: probability distribution (λ = 3.6 ± 0.2, *R*^2^ = 0.999), b) bootstrapping the ML estimate (λ = 3.5 ± 0.1). c) Extrapolation of total blinks vs. threshold yielding *B* = 203 ± 16, for a single dataset, which translates to λ = 3.2 ± 0.3 in this example. Inset: variability of λ measured multiple times in 4 different imaging chambers.

An alternative approach is to use Equation (6) to obtain the ML estimate of λ. In addition, we apply a simple bootstrapping algorithm to the data in order to obtain an empirical measure of the error on that estimate (Figure 2b)^[25]^. This involves randomly sampling 1059 grids from our data set (N=1059), but with replacement after each pick, calculating the ML estimate for the sampled distribution, and reiterating 1000 times. At this low threshold, the analysis yields λ = 3.5 ± 0.1, in agreement with the fit to the cumulative distribution.

We have not yet justified our choice of an intensity threshold. In fact, the value we found for λ will change if we choose a different threshold since this choice will alter the number of blinks detected. To account for this, we instead consider the detected number of blinks at a series of thresholds (Figure 2c). As explained in the previous section, we expect a region where the number of blinks decreases approximately linearly with threshold, which is evident in the figure. We then extrapolate this linear region (dashed line) to zero-intensity threshold to recover blinks missed within the noise. In Figure 2c, we have again performed a bootstrap analysis, but this time on the number of blinks and at each threshold value, to obtain a measure of the error as quantified by the standard deviation of the resulting distribution. We extrapolate this as well (dotted lines), under the assumption that a linear extrapolation is sufficient, to obtain the error at zero threshold. Applying Equation (6) to the extrapolated value *B* = 203 ± 16, we find the ML estimate at the zero-intensity threshold to be λ = 3.2 ± 0.3, where the error bars result from propagating the error in *B* with Equation (6) (see *Supporting Information*).

To obtain Figures 2a and 2b, we pooled data from multiple realizations of the same calibration experiment obtained in several sample imaging chambers. The extrapolation illustrated in Figure 2c is obtained from a single one of these data sets with *N* = 78 dyes. If we repeat this extrapolation for the rest of our datasets, then average the results, we find a final, pooled calibration value of λ = 3.6 ± 0.8. While our example extrapolation in Figure 2c agrees well with the pooled result, a closer look at the data sets reveals a degree of variability in λ. This is displayed in the insert to Figure 2c (for a table of values see *Supporting Information*). What we seem to be observing is the effect of slight variations in buffer conditions on the dye photophysics, which may be due to diffusion-limited reactions between the thiol and the dyes or slight variations in oxygen concentration. Improved buffer conditions^[26]^ or self-healing dyes^[27;28]^ could mitigate this variability, but the message is clearly that one must be cautious both in performing and applying this calibration to a molecular counting experiment.

### 3.2 Calibration of *θ*

We next move on to the target nanostructures used to benchmark our approach. These consist of DNA origami grids with *h* =4 binding sites on each grid complementary to an Alexa-647 labeled ssDNA probe. One utility of these grids is that we can calibrate for the fractional occupancy *θ* directly by imaging since the labeling fluorophores were intentionally spaced within a diffraction limited area, but individually resolvable with SMLM.

In Figure 3a we have arranged a sample of SMLM reconstructions of DNA origami grids, which are randomly distributed on a coverslip, into a lattice. The number of dye labels on each grid was automatically acquired by applying k-means clustering to the localizations. The center-of-mass of each cluster was identified as the spatial coordinate for each fluorescent label, and the resulting spacing of the labels was then compared to the designed template dimensions (see *Supporting Information*). Grids displaying dyes closer than 25 nm apart or further than 100 nm were discarded. In this way, we could filter out poorly imaged grids and detect if there were multiple grids within a diffraction limited spot. A histogram of the number of dyes per grid is shown in Figure 3 with a fit to a binomial distribution. From inspection of our sample population of *N* = 640 surface mounted grids, we find a labeling efficiency of *θ* = 0.51 ± 0.02.

**Figure 3.**
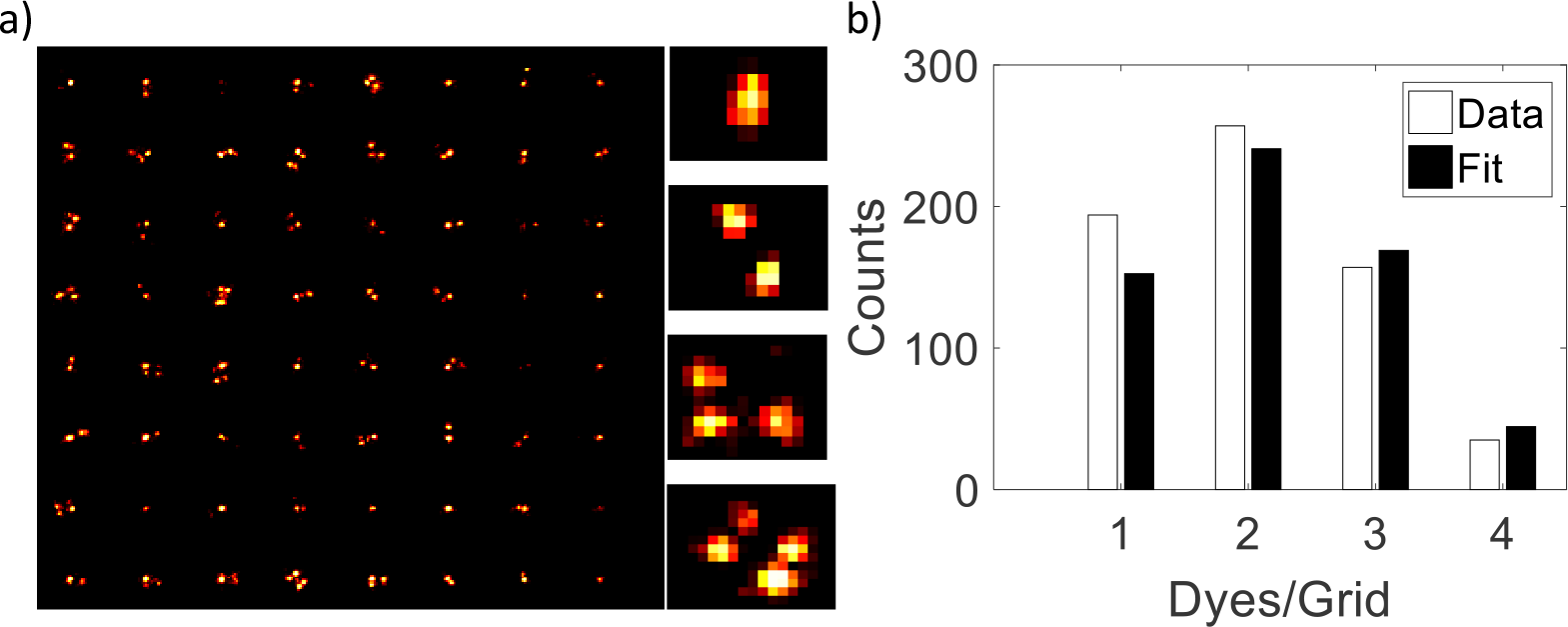
a) SMLM reconstructions of DNA origami grids (N = 640) labeled with up to four Alexa-647 dyes. Enlarged images of a typical grid with 1,2,3 and 4 labels are shown. b) Histogram of the number of dyes per grid with a binomial fit that yields *θ* = 0.51 ± 0.02.

### 3.3 Stoichiometry Analysis

If each fluorescent label corresponds to a single subunit of a complex, a measure of the fractional occupancy *θ* could be interpreted as a quantification of the stoichiometry of that complex. Equation (5), therefore, provides a single-molecule approach to inferring stoichiometry that is an alternative to the traditional method of analyzing photobleaching steps^[29;30;31]^. Single-molecule photobleaching relies upon resolving step-wise changes in intensity as individual, fluorescent labels photobleach. The distribution in the observed number of steps is generated from the data and fit to the appropriate model, which is often a binomial distribution^[32;33]^, yielding a quantification of the stoichiometry. The limited dynamic range of this approach typically restricts its use to cases where there are roughly < 10 fluorophores per diffraction-limited spot, which is a relatively dilute regime for SMLM measurements. While recent advances in step-counting now report that the analysis can be extended well beyond 10 fluorophores^[34;30;35]^, when combined with spatial imaging, the real utility of Equation (5) is it can quantify the stoichiometry of molecular complexes that may not be individually resolvable by diffraction limited techniques.

Here we require knowledge of the characteristic number of blinks λ and *a priori* knowledge of the number of molecular complexes *M*. For the characteristic number of blinks, we will use the pooled, calibrated value found in Section 3.1 (λ = 3.6 ± 0.8). However, because of the sample-to-sample variation we saw in λ, we additionally perform a self-consistent check of each of new dataset analyzed here. This involves using Equation (6) to calculate the ML estimate of λ for each dataset, with *N* equal to the number of single dyes resolved by SMLM within that set, and accepting that set if λ is found to be within one standard deviation of the calibration.

Once again, we consider a dilute concentration of Alexa-647 labeled DNA origami grids, with a maximum of 4 labels each, such that no two grids are within a diffraction-limited spot. In this way, each bright spot we image should correspond to a single complex. The insert in Figure 4 shows a bootstrapping analysis of this data obtained at a low threshold (~ 790 photons). However, a more accurate estimate accounts for blinks missed due to the intensity threshold, so we extrapolate Equation (5) to zero threshold (Figure 4a, main figure). Once again, a bootstrap analysis is performed on the number of blinks at each threshold, and propagated to obtain the error bars shown in the figure (see *Supporting Information*). A linear extrapolation to zero threshold is then made of the estimate (dashed line) and its associated error (dotted lines). For a data set of 49 target complexes, we obtain a fractional occupancy of *θ* = 0.46 ± 0.10, which agrees with the value (*θ* = 0.51 ± 0.02) obtained from directly quantifying the occupancy from the super-resolved image reconstructions. We repeated this analysis for 2 additional datasets with the final ML estimates for the fractional occupancy displayed in Figure 4b (*θ* = 0.46 ± 0.10, 0.51 ± 0.11, and 0.44 ± 0.09).

**Figure 4.**
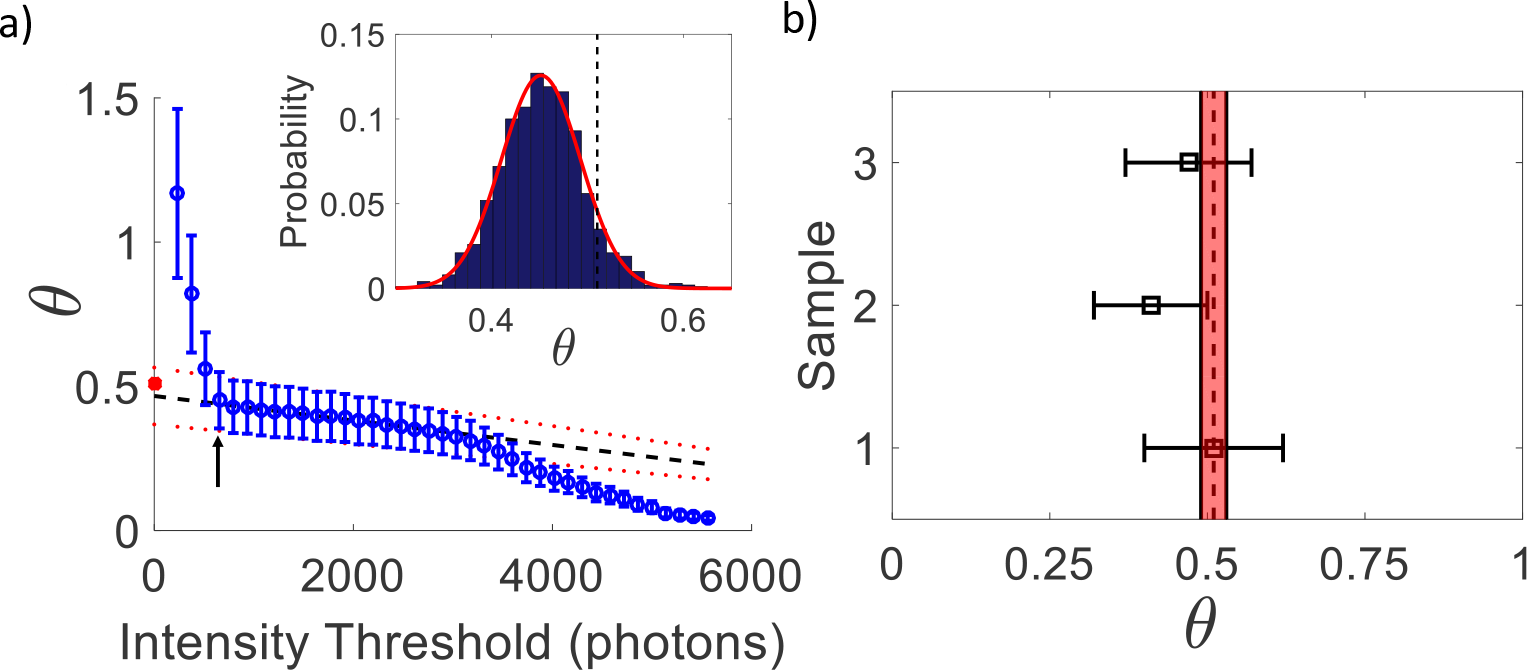
a) ML estimate of the fractional occupancy 0.47 ± 0.10 compared to the result from direct imaging 0.51 ± 0.02 (circle). Dashed (dotted) lines are a linear extrapolation of the mean (error). Insert: histogram of bootstrapping at a low intensity threshold of 790 photons (indicated by arrow in main figure) yielding: *θ* = 0.45±0.04 (dashed line). b) Fractional occupancy obtained from 3 data sets (top to bottom): 0.46 ± 0.10, 0.44 ± 0.09, 0.51 ± 0.11. Vertical dashed (solid) lines indicate the mean (error) from imaging.

### 3.4 Molecular Counting Benchmark

To benchmark our counting method, we again consider a dilute concentration of DNA origami grids such that no two grids are within a diffraction-limited spot, but each bright spot we image now represents a single, binomially labeled ‘molecule’. The target molecules are once again labeled with a maximum of four Alexa-647 dyes. We then compare the number of molecules imaged by SMLM to the number extracted from directly counting blinks. The fractional occupancy of dyes on these target molecules was measured in Section 3.2 and found to be *θ* = 0.51 ± 0.02. However, we will again need to be cautious in calibrating for the characteristic number of blinks λ due to the high degree of sample-to-sample variability observed with single-dye labeled DNA grids in Section 3.1. Here we obtain a self-consistent measure of λ for each dataset by first analyzing a subset of the molecules and applying Equation (6), with the substitution *N* → *Mθh*, to obtain an ML estimate for λ.

With both λ and *θ* in hand, we then evaluate our method’s performance at counting varying numbers of target molecules randomly chosen from the remaining population. Figure 5 displays results from 3 distinct datasets. The characteristic number of blinks for the first set (Figure 5a) was found to be λ_1_ = 3.1 ± 0.3, which is similar to the mean value obtained in Section 3.1 from our pooled, single-dye data (λ = 3.4 ± 0.4). However, we also show results from two additional datasets (Figures 5b,c) where our self-consistent check yielded λ_2_ = 2.0 ± 0.2 and λ_3_ = 7.4 ± 0.7, respectively, which deviates significantly from the pooled, single-dye calibration.

**Figure 5.**
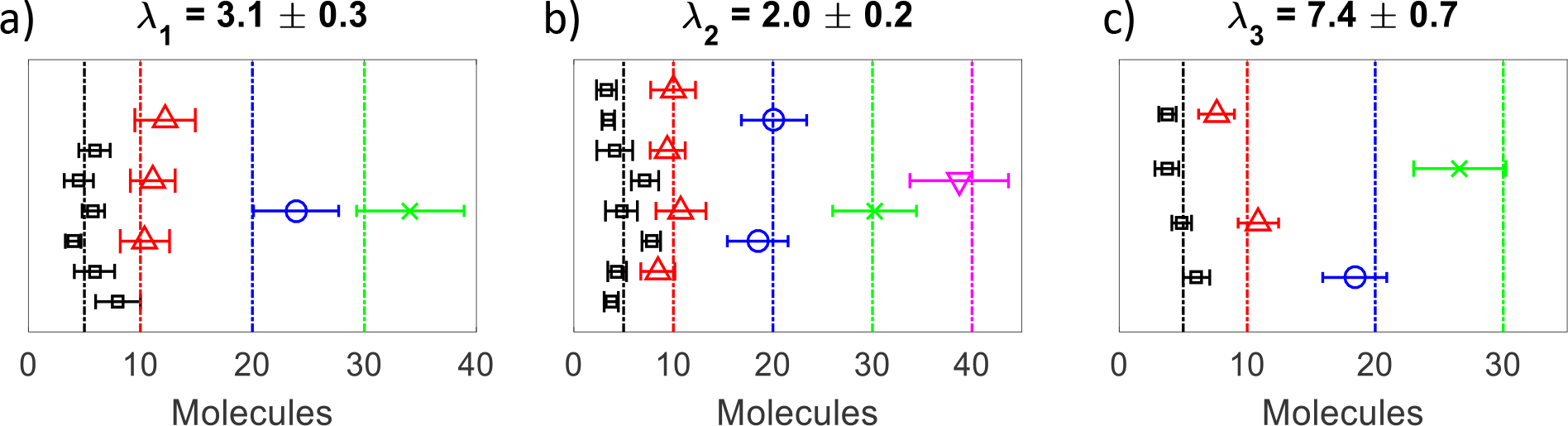
a) From left to right for λ_1_ = 3.1 ± 0.3: target 5, 10, 20, and 30 molecules, b) λ_2_ = 2.0 ± 0.2: target 5, 10, 20, 30 and 40 molecules, c) λ_3_ = 7.4 ± 0.7: target 5, 10, and 20 and 30 molecules.

Results from our molecular counting benchmark experiments, for a range of target molecule number, are shown in Figure 5. To obtain these estimates on the molecular count, and their associated errors, we again performed a threshold analysis, this time on Equation (3). An ML estimate on molecule number was obtained at each threshold value and linearly extrapolated to zero threshold yielding the data points in Figure 5. Likewise, a bootstrap analysis was performed on the number of blinks, at each threshold, and the result was propagated with Equation (3) (see *Supporting Information*) to obtain an empirical measure of the associated error in the estimate of molecule number. The error was then extrapolated to zero threshold to yield the error bars in Figure 5. Examples of the threshold analysis for these two datasets is explicitly provided in the *Supporting Information*.

In Figure 5a, we calibrated λ_1_ = 3.1 ± 0.3 with roughly half of the molecules imaged in the dataset. We then randomly selected from the remaining molecules to generate sets of 5, 10, 20, and 30 molecules. With all target molecules chosen to be unique within a given set, we were able to analyze more instances of smaller molecule number (for example, a single instance of 30 molecules compared to 6 instances of 5 molecules). Figure 5b is a similar analysis on a dataset with λ_2_ = 2.0 ± 0.2. Due to a larger population size, here the data is split into sets of 5, 10, 20, 30, and 40 molecules. Likewise, Figure 5c is for a dataset with significantly larger λ_3_ = 7.4 ± 0.7, which could only be split into sets of 5, 10, and 20 molecules.

The agreement between the ML estimate and the actual number of target molecules highlights the importance of a proper calibration as the pooled, single-dye calibration would not have yielded accurate results for two out of the three data sets. We should also mention that throughout our analyses, we have used bootstrapping to get an empirical measure of the uncertainty in our estimate of molecule number. As an alternative, Equation (5) provides a theoretical relation for this uncertainty. While the two approaches show a reasonable level of agreement, Equation (5) tends to yield higher estimates of the error than the bootstrap prediction, especially for lower numbers of target molecules (see *Supporting Information*). This is not surprising since the accuracy of bootstrapping decreases when the distribution being subsampled is small (i.e., low target molecule numbers), while the theoretical estimate of the error assumes the number of molecules *M* ≫ 1.

### 3.5 Extending the Dynamic Range

Since single-molecule photobleaching is typically limited to counting 8-10 molecules per diffraction-limited spot, we should consider the dynamic range of molecular counting with SMLM, which is reached when blinks from distinct fluorophores begin to temporally overlap. We can set an optimistic theoretical upper bound for the dynamic range based on the duty cycle (DC), as the number of fluorophores that can be detected per diffraction-limited spot varies inversely with the duty cycle, *N*_*f*_ = 1/*DC* (see *Supporting Information*). For a typical SMLM measurement, such as those presented here, the duty cycle is often targeted to be on the order of 10^−4^ or less. Note, however, that if all dyes are initially in the on-state, a period of illumination is required before a sufficient number photoswitch such that the density is dilute enough for single-molecule measurements, which is when the duty cycle is often quoted. During this initial period, some dyes will begin to photoswitch and even photobleach with the duty cycle still relatively high. If the population is significant, it will clearly effect the dynamic range. However, even if we provide for a wide margin of caution, the dynamic range should still be 1-2 orders of magnitude larger than the conventional, photobleaching approach. The photoswitchable properties of organic dyes, such as the duty-cycle, number of switching cycles, and photon yield, are largely determined by the buffer conditions. In particular, the primary thiols typically employed to drive photoswitching in the cyanine dye Alexa-647 are *β*-mercaptoethanol (*β*ME) or mercaptoethylamine (MEA). While both *β*ME and MEA will induce Alexa-647 to photoswitch, the choice is a trade-off between photon yield and duty cycle^[36]^. MEA will generally result in a lower duty cycle than *β*ME, but each photoswitching cycle, or blink, will yield less photons.

In the previous sections, we have performed all experiments in an imaging buffer containing 10 mM MEA. While lower concentrations often yield erratic behavior where not all dyes consistently photoswitch, higher concentrations can lead to a lower duty cycle, but with a diminished number of photons per blink. Although the decreased duty cycle should extend the dynamic range, the lower photon yield is a problem. In Section 2.3, we defined the middle-distribution as the distribution in intensities of those frames where the dye is on in the neighbouring frame both before and after the considered frame. The middle distribution serves as a measure of both the magnitude and steadiness of photon emission by the dye. For a dye emitting with a constant intensity, this distribution would simply be a spike at the full frame intensity; however, more complicated emission dynamics can broaden and shift this distribuiton. In Figure 1 we illustrated how a shift toward lower intensities in the middle distribution can conceal the linear region in our threshold analysis on the number of blinks. Ideally, we would like to reduce the duty cycle while preventing the middle distribution from shifting to lower intensities.

We now consider 3 buffer conditions: 10 mM MEA, 50 mM MEA, and 10 mM MEA with 50 mM βME, applied to single, Alexa-647 labeled DNA origami grids. For these buffers, the measured duty cycles were *DC* = 1.4 × 10^−4^, 0.5 × 10^−4^ and 1.0 × 10^−4^, respectively (see *Supporting Information*). Increasing the MEA concentration to 50 mM significantly lowered the duty cycle while maintaining a 10 mM MEA concentration and incorporating 50 mM *β*ME also lowered the duty cycle, if to a lesser degree. However, if we compare the middle distribution between 10 mM and 50 mM MEA, we see that at 50 mM MEA the distribution shifts toward lower intensities (Figures 6a,b) all but concealing the linear region previously observed in the threshold dependence of the number of blinks (Figure 6c). If we now consider the mixture of 10 mM MEA with 50 mM *β*ME, not only can we reduce the duty cycle, but the middle distribution shifts to even higher intensities than in a buffer of 10 mM MEA alone (Figures 6a,b). Moreover, increased thiol concentrations tend to reduce the variability in photon emission leading to a more peaked middle distribution (Figure 6b).

**Figure 6.**
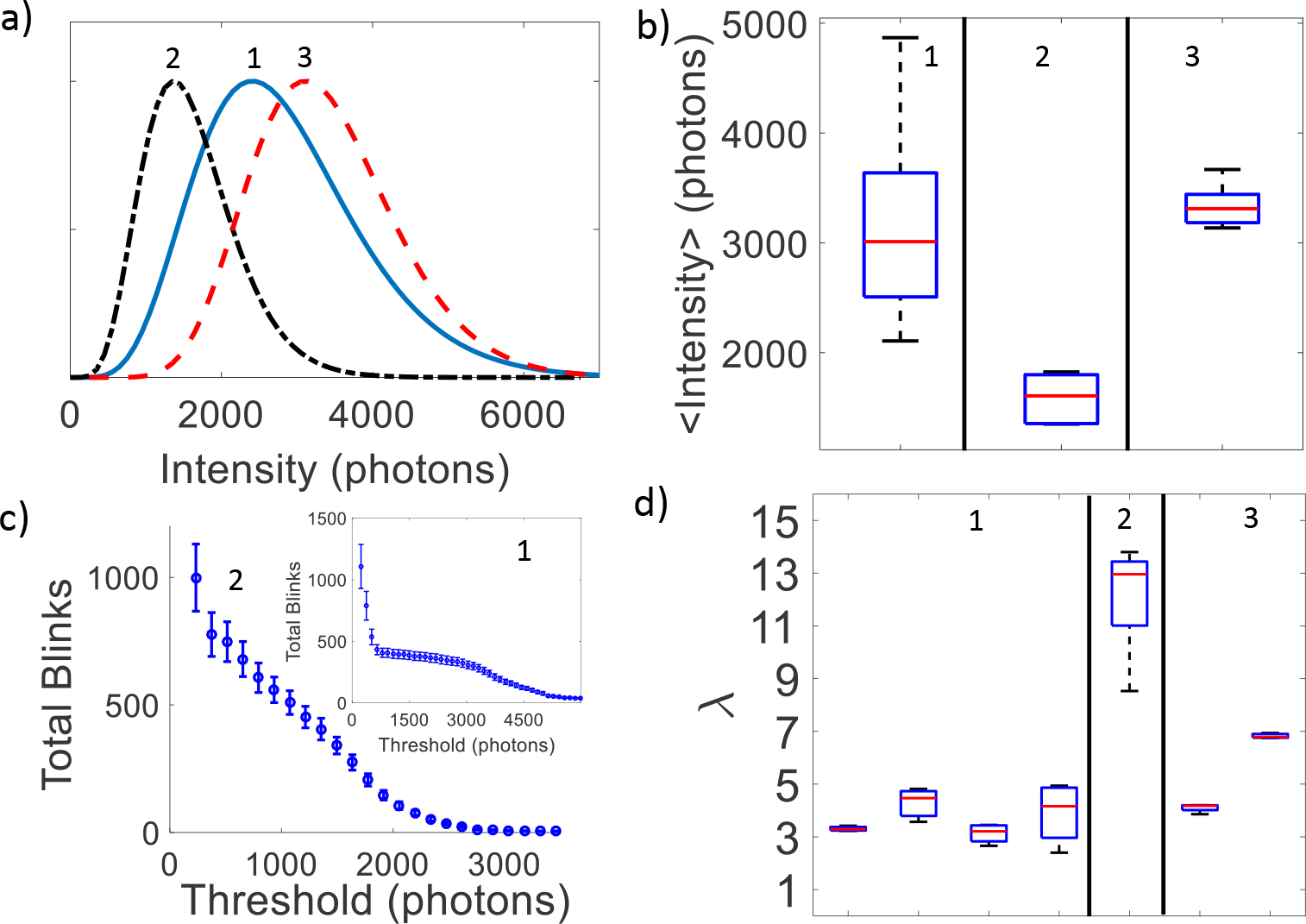
Comparison of 3 buffer conditions: 10 mM MEA (1), 50 mM MEA (2), 10 mM MEA & 50 mM *β*ME (3). a) Middle-distribution intensities. b) Number of blinks as a function of threshold for 50 mM MEA (insert: same at 10mM MEA). c) Variation of mean intensity. d) Variation of characteristic number of blinks λ. Note, λ was obtained at low threshold for 50 mM MEA since a zero threshold extrapolation could not be performed.

Despite our best efforts at reproducibility, for each buffer condition tested, we saw significant variation in our calibration of the characteristic number of blinks (λ) between sample chambers (Figure 6d). Moreover, while some variation was observed over multiple measurements within a single chamber, increasing the concentration of MEA tended to have the detrimental effect of greatly increasing this variability. However, upon supplementing 10 mM MEA with 50 mM *β*ME, this in sample variability was significantly reduced (although, the sample-to-sample variation remained). Once again, interactions between the primary thiols and the dyes may simply be diffusion limited, which could be exacerbated by any fluctuations in residual oxygen interacting with MEA^[37]^, leading to the observed sample-to-sample variation.

## 4. Conclusion

Here we have provided a practical guide to implementing the SMLM counting technique previously developed by Nino *et al.* ^[21]^, with a focus on employing photoswitchable organic dyes. We have shown that, under carefully controlled conditions, this technique can provide an accurate quantification of molecular abundance and stoichiometry within diffraction limited volumes, and should display a dynamic range rivalling more conventional single-molecule techniques.

Perhaps the greatest challenge in implementing this technique is accurately calibrating the single-dye photophysics by measuring the characteristic number of blinks λ. In fact, even in the well controlled *in vitro* system we used to benchmark our method, we had difficulty in obtaining a consistent measure of λ across sample preparations. Instead, for the present experiments, we resorted to extracting λ self-consistently from a fraction of the molecules imaged in each SMLM experiment, and either used this self-consistent value to confirm the calibration or employed it instead. We could do this because our target molecules (here, DNA origami grids) were themselves resolvable via SMLM, so we could already count molecules or quantify the stoichiometry purely through imaging, which is typically not the case.

One could, however, imagine introducing a control population of resolvable targets that would be used to calibrate λ. Alternatively, this method could be applied in systems where protein dimerization or higher order oligomerization is inducible, which is common throughout cell biology^[38]^ (consider transmembrane proteins such as G-protein coupled receptors (GPCRs)^[39]^ or ATP-binding cassette (ABC) transporters^[40]^). Here the calibration of λ would be performed prior to the induction of oligomerization, where the individual proteins could be resolved.

Still, for this technique to be adopted as a universal approach to molecular counting, it is necessary to reduce the sample-to-sample variability in λ. While improvements in sample preparation might be possible, a more robust strategy would be to reduce the environmental sensitivity of the dyes, for instance, by developing better imaging buffers. We found that supplementing our 10mM MEA buffer with 50 mM *β*ME yielded more stable calibrations within a sample prep. But other enhanced imaging buffers have been proposed that might translate to enhanced counting buffers. For instance, addition of the polyunsaturated hydrocarbon cyclooctatetraene (COT) to the SMLM imaging buffer has been reported to result in a 3- to 4-fold improvement in photon yield^[26]^. In the presence of COT, Alexa-647 was observed to maintain an enhanced stability, which is thought to be through quenching of the triplet state^[41]^, while maintaining its photoswitching characteristics, such as a low duty cycle. This was not observed with other triplet state quenchers, such as Trolox, that interfered with photoswitching. Of course, a concern when tuning the photoswitching with buffer composition, especially in cells, is that the local concentration of buffer components cannot be conserved throughout a sample leading to spatial variation in the dye photophysics. A better route, perhaps, is through employing self-healing dyes. For example, TrisNTA-Alexa647 (NTA, N-nitrilotriacetic acid), which is Alexa-647 linked to a Ni(II) triplet state quencher, shows both improved photostability as well as a 25-fold enhancement in photon emission per blink^[27]^.

Molecular counting via SMLM holds significant promise for optical ‘-omics’ technologies due to its extreme sensitivity, high signal-to-noise ratio, low sample consumption, and rapid analysis time. And while current proteomic technologies might be unable to detect low abundance proteins or local protein concentrations, this information is directly accessible to SMLM counting. Moreover, robust SMLM counting methods might eliminate the need for an amplification stage prior to sequencing of DNA or RNA, which would significantly increase the accuracy and reliability of single-cell genomic analyses. While currently confined to the research laboratory, as these techniques mature they should significantly impact ‘-omics’ related biotechnology, from basic research applications to drug-screening and medical diagnostics.

## 5. Experimental Section

### Flow Chambers

Glass coverslips (0.17 mm thickness) and microscopy slides were washed in 3 M KOH in a sonicator bath for 15 min then rinsed 3 times with distilled water. They were subsequently sonicated in 95% EtOH for 15 min and again rinsed 3 times with distilled water. The cleaned coverslips were incubated in 10% poly-l-lysine (PLL) solution overnight. Gold nanoparticles (40 nm diameter, Sigma Aldrich) were used as fiducial markers for drift correction. The nanoparticles were diluted in water by a factor of 20 (making a 5% solution), which was used to cover the entire coverslip and incubated at room temperature for 25 min. The coverslips were then rinsed with distilled water and air dried. Flow chambers were constructed with double-sided tape to make flow channels of roughly 20 *μ*l in volume.

### Assembly and Mounting of DNA Origami Grids

DNA origami grids were assembled following^[42]^ with external labeling. We found 15 mM MgCl_2_ to be the optimal concentration for well-formed grids. Two variants of the grids were constructed (see *Supporting Information*). To perform the photophysics calibration experiments, we built grids with a single docking strand in the middle of the grid to which a complimentary probe end-labeled with a single Alexa-647 dye could bind. To perform the counting experiments, we built grids with a docking strand at each of the corners of the grid, simulating a molecule with 4 possible binding sites. The advantage of spacing out the labeling sites is that we can obtain a measure of the labeling efficiency directly from the super-resolved images of the grids.

To mount the DNA origami grids to the surface, we first flushed 50 *μ*l of 0.5 mg/ml BSA-Biotin into each channel and incubated at room temperature for 10 min. The channels were then washed with 100 *μ*l of PBS (pH 7.4, filtered through a 0.2 *μ*m filter). 50 *μ*L of 10% BSA solution was twice added to each channel to block any remaining exposed PLL. Each round of BSA was incubated for 10 minutes at room temperature and then washed with 100 *μ*l PBS. This was followed by adding 50 *μ*l of 0.5 mg/ml strepavidin to each channel and incubating at room temperature for 10 minutes, followed by a 100 *μ*L wash with PBS buffer. DNA origami grids were diluted 500× to 1000× in folding buffer (5mM Tris, 50mM NaCl, 1mM EDTA, and 15 mM of MgCl_2_). 50 *μ*l of the DNA origami solution was added to each channel and incubated for 10 minutes. The chamber was then washed with a 1× folding buffer solution.

### Single-Molecule Localization Microscopy

The microscopy was performed on a home-built setup based upon an Olympus IX-81 inverted microscope with a 60x, NA=1.49 oil-immersion TIRF objective (Olympus, APON 60XOTIRF). All associated instrumentation and data acquisition was controlled via *μ*Manager^[43]^. We continuously excited the sample with a 637 nm laser (World Star Tech) at 2.4 kW*/*cm^2^ to induce the dyes to photoswitch. Images were captured on an EMCCD camera (iXion3, Andor Technology) with an EM gain of 274. The effective pixel-size of the camera was 117 nm after magnification by an additional telescope placed just before the camera.

Our imaging buffer (pH=8.0) consisted of a protocatechuic acid/protocatechuic dioxygenase (PCA/PCD) oxygen scavenging system (13 mM PCA, 50 nM PCD) in folding buffer with the thiolating compounds MEA or BME added to induce photoswitching. The imaging buffer was introduced into the sample chamber and allowed to incubate for at least 30 minutes prior to all measurements to allow sufficient time for oxygen removal. We found that the molecular counting experiments were affected by the choice and concentrations of MEA and BME. These effects are discussed in Section 3.5, but unless otherwise noted, the microscopy was performed at 10 mM MEA.

### Drift Correction

We correct for axial drift in real time with an autofocus system (CRISPR, ASI). Lateral drift is corrected post-acquisition by tracking of 40 nm gold fiducial markers. We crop a 20×20 pixel region around the fiducial markers and find their central coordinates at each frame by a 2D Gaussian fit. As the gold nanoparticles are much brighter than the dyes, we set a high intensity threshold to ensure the surrounding dyes are not detected. We then clean up the drift time-series by applying a running median filter with a window size of 650 frames. Finally, we take the median of the total drift over several gold nanoparticles to eliminate spurious drift from any one particle (see *Supporting Information*).

### Data Analysis

To accurately determine the number of blinks, we need to carefully construct a table of localizations. Here, we construct the localization table using rapidSTORM, a popular open-source localization program for low-density data sets^[44]^. RapidSTORM detects spots of interest as the maximums of a smoothed image and fits a 2D Gaussian PSF by the Levenberg-Marquardt least squares method.

To identify DNA origami grids, bright spots in the initial frame, after turning on the excitation laser, are labeled as candidates and further analyzed. Fiducial markers and identified spots that displayed an excessive number of localizations were filtered out based on the distribution of the number of localizations per fluorophore (see *Supporting Information*). Spots that were within one pixel of each other or five pixels of a gold nanoparticle, within 3 pixels from the edge of the ROI, or had drifted out of the ROI during acquisition, were also discarded. Note, as we do not count the initial burst as a blink, we discard localizations coming from the initial burst.

Once the molecules are identified, to then count blinks, we detect the number of localization events happening in consecutive frames within a certain spatial radius from the initial molecule coordinates. We found that rapidSTORM sometimes misses localizations within a few frames during each burst. Without correction, this would artificially increase the detected number of blinks. Rather than assign a global temporal radius to bunch localizations, as is typically done, we reanalyze the missing frames directly on the stack by trying to fit a 2D Gaussian to the spot where the fluorophore had last been localized. If the Gaussian fit parameters (intensity and width *σ*_*x*_, *σ*_*y*_) fall within the bounds of other successfully detected localizations, we effectively fill in the hole then bunch all the consecutive localizations into one burst. This procedure is especially important when we determine the number of blinks as a function of intensity threshold.

## Supporting Information

Supporting information is available.

## Acknowledgements

D.N. thanks Anna Shahmuradyan for helpful discussions. This work was funded by the Natural Sciences and Engineering Research Council of Canada [J.N.M., D.N., D.D.] and an Early Researcher Award from the Ministry of Research and Innovation [J.N.M.].

