## Supporting Information for "Nanoscopic Stoichiometry and Single-Molecule Counting"

#### Contents

1 **Drift Correction** 2

2 **DNA Origami** 3

3 **On-Times Distribution and Duty Cycle** 4

4 **Theoretical Upper Bound on the Dynamic Range** 5

5 **Total Localizations per Fluorophore** 6

6 **Reproducibility of $\lambda$** 7

7 **Poisson Labeling Statistics** 8

8 **Error Propagation** 9

9 **Counting Molecules** 10

10 **Uncertainties** 11

### 1 Drift Correction

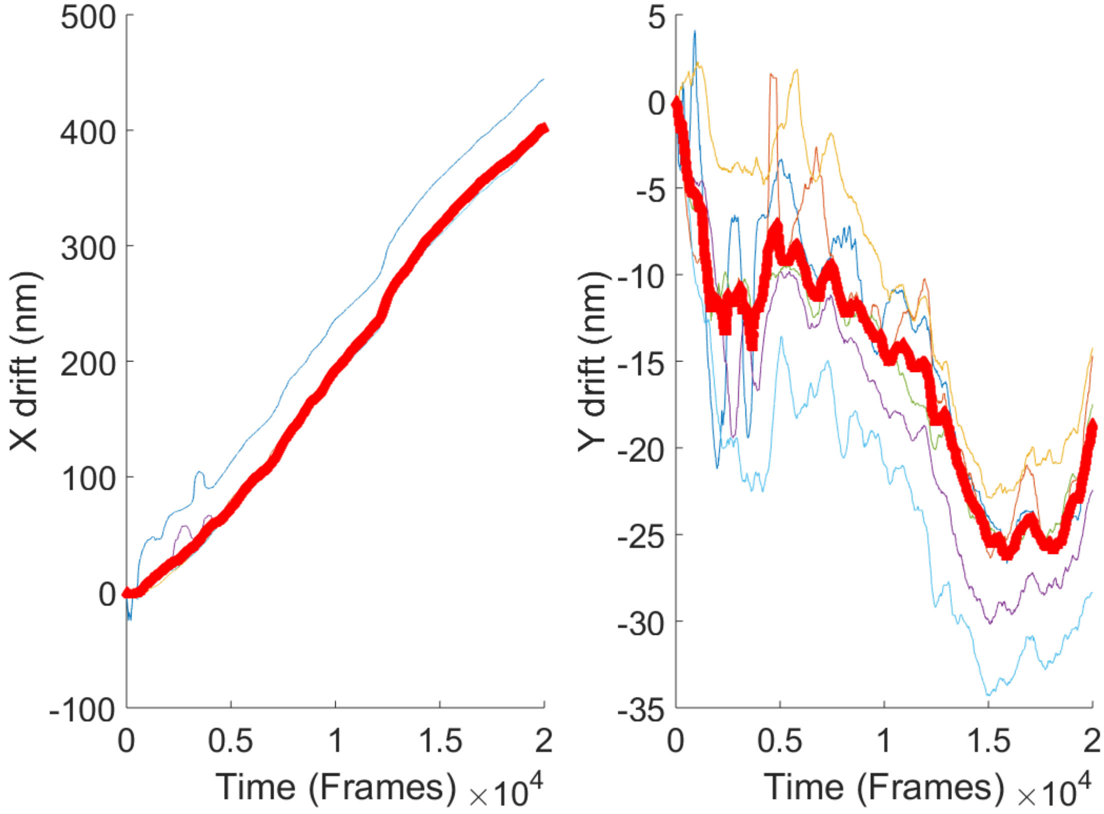


Figure 1: Example time traces for multiple fiducial markers. Thick red line is the median of all the trajectories.

#### 2 DNA Origami

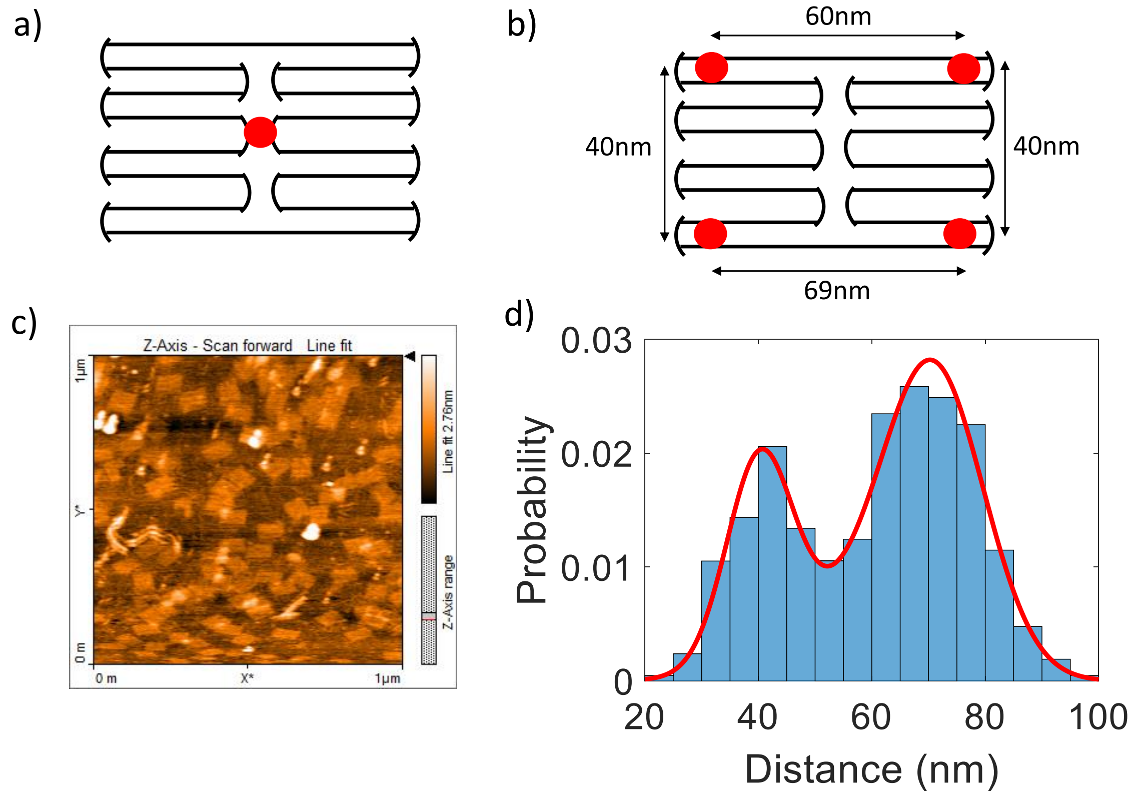


Figure 2: a) Design of DNA Origami grids with single binding site for a dye in the middle. b) Design of DNA Origami grid with 4 possible binding sites. c) Atomic force microscopy image of DNA Origami grids, confirming formation of grids. d) Histogram of distances between dyes in DNA Origami with 4 possible binding sites (n = 418 ). Red curve is a Gaussian mixture model fit with 3 components, yielding mean distances of 40.3 nm, 61.2 nm and 71.2 nm, as designed.

### 3 On-Times Distribution and Duty Cycle

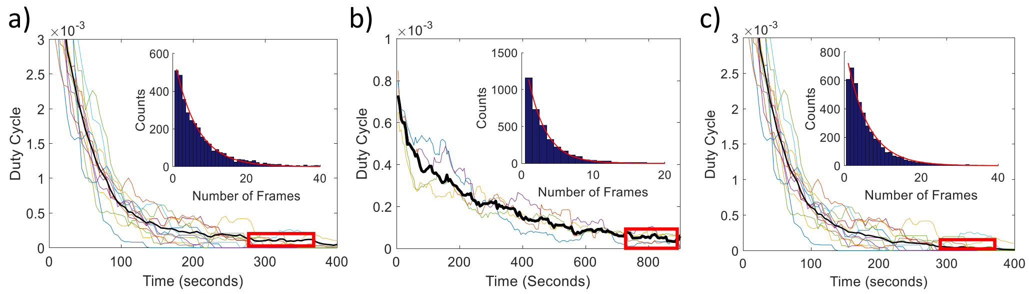


Figure 3: Duty cycle (main figures) and ON-Time distribution (inserts). a) 10mM MEA: DC: $1.4 \times 10^{-4}$ , $T_{ON} = (6.1 \pm 0.2)$ frames. b) 50mM MEA, DC: $5.0 \times 10^{-5}$ , $T_{ON} = (2.5 \pm 0.1)$ frames c) 10mM MEA + 50mM $\beta$ME: DC: $1.0 \times 10^{-4}$ , $T_{ON} = (5.6 \pm 0.3)$ frames, all taken at an intensity threshold of 700 photons, each frame is 50ms. Duty cycle was measured using a 50s sliding window.

### 4 Theoretical Upper Bound on the Dynamic Range

Defining the duty cycle as $DC = T_{ON}/(T_{ON}+T_{OFF})$, which is the probability of a fluorophore being in the ON state at any given time, the probability that a fluorophore is off is then $p_{OFF} = 1 - DC$. Let $p_N(M)$ be the probability of $M$ fluorophores being on out of a population of N fluorophores. In a diffraction-limited spot, the probability of *no more than* 1 fluorophore being ON at any time is equal to $p_N(1)+p_N(0) = NDC(1-DC)^{N-1}+(1-DC)^N$. Under typical conditions for super-resolved experiments, the DC is usually $10^{-4} - 10^{-3}$, so we can Taylor expand about the DC:

$$p_N(1) + p_N(0) \approx 1 - \frac{N(N-1)}{2}DC^2. \tag{1}$$

So, the probability of *more than one* fluorophore being on in a diffraction-limited spot is:

$$p_N(>1) \approx \frac{N(N-1)}{2}DC^2. \tag{2}$$

For N = 1/DC, the probability of having more than fluorophore on within a diffraction-limited spot is 50%.

### 5 Total Localizations per Fluorophore

To identify fluorophores which behaved erratically, we used the distribution of total number of localizations to remove fluorophores which had too many localizations. This was only needed in the case of 10mM MEA, where some fluorophores blinked an excessive number of times. Data is presented at an intensity threshold of 700 photons for different buffer conditions. Localizations from the initial burst were removed.

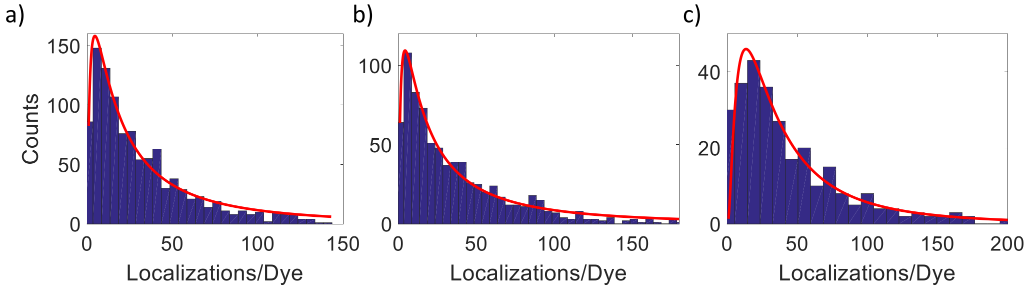


Figure 4: Number of localizations per dye. a) 10 mM MEA b) 10 mM MEA, 50 mM BME c) 50 mM MEA. Red curve is a fit to a log-normal distribution.

#### 6 Reproducibility of $\lambda$

| Date | $\lambda$ | $\lambda$ | $\lambda$ | $\lambda$ |
|---|---|---|---|---|
| Sample 1 | $3.42 \pm 0.64$ | $3.33 \pm 0.65$ | $3.23 \pm 0.37$ | $3.25 \pm 0.32$ |
| Sample 2 | $4.82 \pm 0.57$ | $3.57 \pm 0.43$ | $4.47 \pm 0.47$ | N/A |
| Sample 3 | $2.66 \pm 0.34$ | $3.24 \pm 0.32$ | $3.45 \pm 0.39$ | $2.32 \pm 0.26$ |
| Sample 4 | $4.94 \pm 0.40$ | $4.79 \pm 0.46$ | $3.53 \pm 0.37$ | $2.40 \pm 0.30$ |

Table 1: Values of $\lambda$ associated with Figure 2c (inset) showing the variation within and across experiments.

### 7 Poisson Labeling Statistics

In certain circumstances, such as when considering antibody labeling, a Poisson distribution is a better choice for the labeling distribution. The probability of finding N poisson distributed labels among M molecules is:

$$p(N|M) = \frac{(\eta M)^N e^{-\eta M}}{N!}. \tag{3}$$

Equations (3),(4) and (5) can be extended to account for Poisson labeling statistics simply by taking the limits $\theta \to 0$ and $h \to \infty$ such that $h\theta \to \eta$. This yields the following relations:

$$\tilde{\mu}_M = \frac{B(1 - e^{-1/\lambda})}{\eta}, \tag{4}$$

$$\tilde{\sigma}_M^2 = \frac{\tilde{\mu}_M}{\eta}(1 + e^{-1/\lambda}), \tag{5}$$

and

$$\eta_{ML} = \frac{B(1 - e^{-1/\lambda})}{M}. \tag{6}$$

### 8 Error Propagation

The MLE for the number of molecules given the number of blinks depends on experimentally determined parameters. As such, to find the experiment error on the MLE, we propagate the error:

$$
\begin{aligned}
\delta_{\mu_M}^2 &= \left(\frac{\partial \mu_M}{\partial B}\right)^2 \delta_B^2 + \left(\frac{\partial \mu_M}{\partial \lambda}\right)^2 \delta_\lambda^2 + \left(\frac{\partial \mu_M}{\partial \theta}\right)^2 \delta_\theta^2 \\
&= \left(\frac{1 - e^{-1/\lambda}}{\theta h}\right)^2 \delta_B^2 + \left(\frac{B e^{-1/\lambda}}{\theta h \lambda^2}\right)^2 \delta_\lambda^2 + \left(\frac{B(1 - e^{-1/\lambda})}{\theta^2 h}\right)^2 \delta_\theta^2.
\end{aligned} \tag{7}
$$

Similarly for the characteristic number of blinks, we have

$$
\delta_{\lambda_{ML}}^2 = \left(\frac{\partial \lambda_{ML}}{\partial B}\right)^2 \delta_B^2 = \left(\frac{\lambda_{ML}^2 N}{B(N - B)}\right)^2 \delta_B^2. \tag{8}
$$

For the calibration experiments, we use a generalization of the MLE for the characteristic number of blinks:

$$
\frac{1}{\lambda_M} = -\ln\left(1 - \frac{h\theta M}{B}\right). \tag{9}
$$

The propagated error for this:

$$
\begin{aligned}
\delta_{\lambda_M}^2 &= \left(\frac{\partial \lambda_M}{\partial B}\right)^2 \delta_B^2 + \left(\frac{\partial \lambda_M}{\partial \theta}\right)^2 \delta_\theta^2 \\
&= \left(\frac{\lambda_M^2 h\theta M}{B(h\theta M - B)}\right)^2 \delta_B^2 + \left(\frac{\lambda_M^2 h M}{B}\right)^2 \delta_\theta^2.
\end{aligned} \tag{10}
$$

For the labeling efficiency:

$$
\begin{aligned}
\delta_{\theta_{ML}}^2 &= \left(\frac{\partial \theta_{ML}}{\partial B}\right)^2 \delta_B^2 + \left(\frac{\partial \theta_{ML}}{\partial \lambda}\right)^2 \delta_\lambda^2 \\
&= \left(\frac{\theta_{ML}}{B}\right)^2 \delta_B^2 + \left(\frac{B e^{-1/\lambda}}{h M \lambda^2}\right)^2 \delta_\lambda^2.
\end{aligned} \tag{11}
$$

#### 9 Counting Molecules

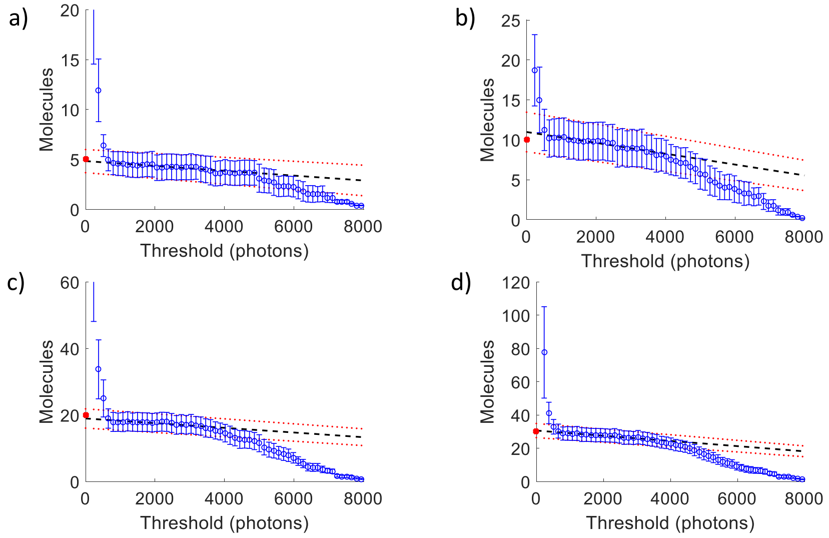


Figure 5: Sample curves of number of estimated molecules as a function of intensity threshold for $\lambda = 2.0 \pm 0.2$. Dashed black line shows linear extrapolation to zero threshold. Dashed red curves show error. Error bars were estimated with bootstrapping.

### 10 Uncertainties

Values associated with Figure 5 of the main text. Target Molecules is the number of randomly chosen molecules. $\tilde{\mu}_M$ is the ML estimate of the number of molecules, $\sigma_{exp}$ is the experimental (bootstrapping) uncertainty associated with the ML estimate, and $\tilde{\sigma}_M$ is the theoretical error calculated from Equation (4) of the main text.

| Target Molecules | $\tilde{\mu}_M$ | $\sigma_{exp}$ | $\tilde{\sigma}_M$ |
|---|---|---|---|
| 30 | 34.1 | 4.8 | 4.5 |
| 20 | 23.9 | 3.8 | 3.8 |
| 10 | 10.4 | 2.2 | 2.5 |
| 10 | 11.1 | 2.0 | 2.5 |
| 10 | 12.2 | 2.7 | 2.7 |
| 5 | 8.0 | 2.0 | 2.2 |
| 5 | 5.9 | 1.8 | 1.9 |
| 5 | 4.0 | 0.7 | 1.5 |
| 5 | 5.8 | 1.0 | 1.9 |
| 5 | 4.5 | 1.3 | 1.6 |
| 5 | 5.9 | 1.4 | 1.9 |

Table 2: $\lambda = 3.1 \pm 0.3$

| Target Molecules | $\tilde{\mu}_M$ | $\sigma_{exp}$ | $\tilde{\sigma}_M$ |
|---|---|---|---|
| 40 | 38.7 | 4.9 | 4.6 |
| 30 | 30.2 | 4.3 | 4.0 |
| 20 | 18.5 | 3.0 | 3.2 |
| 20 | 20.2 | 3.3 | 3.3 |
| 10 | 8.5 | 1.7 | 2.1 |
| 10 | 10.8 | 2.5 | 2.4 |
| 10 | 9.4 | 1.8 | 2.3 |
| 10 | 10.0 | 2.3 | 2.3 |
| 5 | 3.3 | 0.7 | 1.3 |
| 5 | 3.5 | 1.0 | 1.4 |
| 5 | 4.1 | 0.9 | 1.5 |
| 5 | 7.2 | 1.6 | 2.0 |
| 5 | 4.8 | 1.4 | 1.6 |
| 5 | 7.8 | 1.8 | 2.0 |
| 5 | 4.4 | 0.6 | 1.5 |
| 5 | 3.8 | 1.0 | 1.5 |

Table 3: $\lambda = 2.0 \pm 0.2$

| Target Molecules | $\tilde{\mu}_M$ | $\sigma_{exp}$ | $\tilde{\sigma}_M$ |
|---|---|---|---|
| 30 | 26.7 | 3.6 | 4.2 |
| 20 | 18.4 | 2.5 | 3.5 |
| 10 | 7.6 | 1.4 | 2.2 |
| 10 | 10.9 | 1.6 | 2.7 |
| 5 | 3.8 | 0.7 | 1.6 |
| 5 | 3.7 | 0.9 | 1.6 |
| 5 | 4.9 | 0.8 | 1.8 |
| 5 | 6.0 | 1.0 | 2.0 |

Table 4: $\lambda = 7.4 \pm 0.7$
